# Wastewater microorganisms impact microbial diversity and important ecological functions of stream periphyton

**DOI:** 10.1101/2022.04.27.489724

**Authors:** Louis Carles, Simon Wullschleger, Adriano Joss, Rik I.L. Eggen, Kristin Schirmer, Nele Schuwirth, Christian Stamm, Ahmed Tlili

## Abstract

Wastewater treatment plant effluents can impact microbial communities in receiving streams. However, little is known about the role of microorganisms in wastewater as opposed to other wastewater constituents, such as nutrients and micropollutants. We aimed therefore at determining the impact of wastewater microorganisms on the microbial diversity and function of periphyton, key microbial communities in streams. Periphyton was grown in flow-through channels that were continuously alimented with a mixture of stream water and unfiltered or ultra-filtered wastewater. Impacts were assessed on periphyton biomass, activities and tolerance to micropollutants, as well as on microbial diversity. Our results showed that wastewater microorganisms colonized periphyton and modified its community composition, resulting for instance in an increased abundance of Chloroflexi and a decreased abundance of diatoms and green algae. This led to shifts towards heterotrophy, as suggested by the changes in nutrient stoichiometry and the increased mineralization potential of carbon substrates. An increased tolerance towards micropollutants was only found for periphyton exposed to unfiltered wastewater but not to ultra-filtered wastewater, suggesting that wastewater microorganisms were responsible for this increased tolerance. Overall, our results highlight the need to consider the role of wastewater microorganisms when studying potential impacts of wastewater on the receiving water body.

**Environmental implication:** The present study investigates the impact of wastewater microorganisms on periphyton, i.e. communities forming the microbial skin of streambeds. We were able to disentangle specific effects of wastewater microorganisms in the context of the complex wastewater matrix. Indeed, wastewater microorganisms induced strong changes in periphyton community composition and function, suggesting the need to consider wastewater microbial communities as a stressor *per se*, similarly to, e.g., nutrients and micropollutants. Moreover, since periphyton is at the basis of the food web in streams, these changes may have consequences for higher trophic levels.

## 1. Introduction

Wastewater treatment plants (WWTPs) represent a major point source for surface water pollution in urban areas, potentially leading to the degradation of the ecosystem ecological status by affecting the structure and function of aquatic communities in the receiving streams (Gessner and Tlili, 2016; Stamm et al., 2016; Vörösmarty et al., 2010). Effluents from WWTPs typically contain various constituents such as dissolved organic matter, microorganisms and complex mixtures of micropollutants, which are not completely retained by the treatment processes. Due to such chemical and biological complexity, disentangling the specific effects of wastewater constituents on receiving aquatic ecosystem from each other is a key challenge. This requires a study design that allows for controlled interventions, which cannot be easily achieved in natural environments. We therefore constructed a flow-through channel system that mimics the complexity of field conditions while allowing for targeted manipulations, and used stream periphyton as a biological model to explore the impact of WWTP effluents.

Stream periphyton, also referred as aquatic biofilms, is a highly diverse and dynamic community of prokaryotic and micro-eukaryotic organisms that are embedded in an extracellular matrix, attached to the surface of submerged solid substrata. Periphyton plays a crucial role in streams as a basis of aquatic food webs and by contributing to important ecological processes such as primary production and nutrient cycling (Battin et al., 2016). With its high biological diversity and important ecological role, periphyton is widely used as a biological community model to assess effects of biotic and abiotic environmental factors, such as trophic interactions, chemical pollution or eutrophication (Montuelle et al., 2010; Sabater et al., 2007). Several field and mesocosm studies have reported shifts in the microbial diversity of periphyton upon exposure to treated wastewater (Carles et al., 2021; Peng et al., 2018; Romero et al., 2019; Tamminen et al., 2022; Tardy et al., 2021; Tlili et al., 2020, 2017). They have also shown that these shifts were accompanied by changes in important functional traits of the communities, such as tolerance to micropollutants, algal primary production and photosynthesis or bacterial secondary production. Notwithstanding these findings, most of these studies focused either on the overall effluent toxicity, or on specific wastewater constituents, such as nutrients and micropollutants, but not on the microorganisms from the WWTP.

It has been demonstrated that downstream of the WWTP, bacterial community profiles in the water column (i.e. planktonic) resembled a mixture of the upstream and the effluent communities (Mansfeldt et al., 2020; Price et al., 2018). Periphyton, however, has a different microbial lifestyle than planktonic cells: species interactions play a major role in determining microbial colonization dynamics and in shaping the final periphyton community structure. This means that changes in taxa composition in periphyton cannot be predicted based on hydraulic mixing alone. Systematic studies focusing on the invasion of stream periphyton by microorganisms from WWTPs and the resulting consequences for community structure and function are scarce. For instance, Chonova et al. (2019) have shown that 27% of diatom taxa found in the periphyton downstream of urban and hospital WWTPs potentially originated from the wastewater effluents. In a similar vein, Mußmann et al. (2013) showed that, despite the high diversity of bacterial nitrifiers in the wastewater effluents, only two taxa colonized the downstream periphyton. Nevertheless, monitoring the taxonomic profiles of these microorganisms alone is insufficient, as it does not inform about their contribution to changes in community functions.

Given this background, we aimed in this study to understand the specific contribution of the microorganisms that originate from WWTPs to the composition and functions of periphyton communities. To this end, periphyton was grown in flow-through channels that were continuously alimented with stream water mixed with various fractions of unfiltered or ultra-filtered treated urban wastewater. Ultrafiltration of the wastewater was intended to remove more than 99% of microorganisms while leaving dissolved nutrients and micropollutants unaffected. We evaluated the effects of the wastewater on periphyton by measuring a set of functional and biomass endpoints that targeted specifically the heterotrophic and phototrophic components of the community. We also determined the tolerance of periphyton communities towards a micropollutant mixture extracted from passive samplers deployed in the wastewater effluent. Moreover, we performed a detailed analysis of the algal and bacterial community structure via genomic high-throughput sequencing, and a comprehensive quantification of micropollutant concentrations in the water and in periphyton samples.

## 2. Material and methods

### 2.1. Experimental system and design

The experiment was conducted in a flow-through channel system called “Maiandros” that mimics natural conditions in freshwater streams but allows for targeted manipulations. Initially described by Burdon et al. (2020), this system was subsequently equipped with a light system to ensure a photoperiod of 12h light: 12h dark (Carles et al., 2021), as well as with an automated-online monitoring system for parameters such as flow rate, water conductivity and temperature (Desiante et al., 2022). The ultra-filtration (UF) unit added to the system consisted of a membrane module of 50 m^2^ surface and a nominal pore size of 0.4 μm, allowing for the removal of particulates (including microorganisms) from the effluent, with a microorganism removal efficiency of 99.3% (Desiante et al., 2022). The Maiandros system allowed controlled mixing of different waters for comparative experiments in 20 independent flow-through channels (total length of 2.6 m, 0.15 m width and 0.1 m water depth). Each channel is equipped with a paddle wheel providing 0.2 m s^-1^ horizontal flow speed. The channels were continuously fed with a mixture of stream water that was directly pumped from Chriesbach, a small peri-urban stream (47°24’16.7”N 8°36’41.4”E; Dübendorf, Switzerland), and wastewater treated for nitrification and denitrification (Desiante et al., 2022), thereafter called “wastewater”. The 20 channels were randomly assigned to five treatments (n = four independent replicate channels per treatment), corresponding to a nominal proportion of 0% (control), 30% and 80% of either unfiltered (WW) or ultra-filtered (UF) wastewater.

### 2.2. Periphyton colonization and sampling

Periphyton was grown on clean glass slides (210 × 75 × 4 mm) that were placed vertically in the channels (40 glass slides per channel). After 28 days, colonized slides were retrieved and transported to the laboratory for chemical and biological analyses as described by Carles et al. (2021). Briefly, periphyton growing on five glass slides from the same channel was scraped, pooled and suspended in 200□mL of Evian natural water. Fresh suspensions were used for the biological analyses (see section 2.5. below) while the remaining volume was lyophilized before micropollutant analysis and Carbon:Nitrogen:Phosphorus (C:N:P) ratio determination.

### 2.3. Water sampling

Two AttractSPE®Disks SDB-RPS (47 mm diameter, Affinisep, France) with Supor^®^ polyethersulfone (PES) membrane disc filters (47 mm diameter, 0.45 μm pore size, VWR, Switzerland) were installed in two channels of each treatment and in the wastewater effluent to sample polar to semi-polar organic micropollutants according to Moschet et al. (2015). The passive samplers were deployed during twelve days and renewed for a second twelve-day period. These passive samplers were then used for micropollutant analysis. The same type of passive samplers but without PES membrane (i.e. to allow faster and higher accumulation of micropollutants) were also deployed in the wastewater buffer tank; from these, the accumulated micropollutants were extracted and used for the community tolerance bioassays described in section 2.5.2 below (Tlili et al., 2017).

In order to characterize the microbial community in all water sources by next generation sequencing, three composite water samples were taken weekly from the stream water as well as from the unfiltered and ultra-filtered wastewater during 24-hours with an automated water sampler (Maxx, TP5 C Aktiv, Germany). The water samples were automatically taken (50 mL every 30 minutes), kept at 4°C, pooled together and immediately filtrated on Supor® polyethersulfone membrane disc filters with 0.2 μm pore size (Pall Corporation, USA). The filters were stored at −80 °C prior to DNA extraction (see section 2.5.3 below).

### 2.4. Micropollutant analyses

A total of 54 organic micropollutants, consisting of 22 pesticides, 25 pharmaceuticals, 4 artificial sweeteners, 2 corrosion inhibitors and caffeine were analysed in extracts from the passive samplers and in periphyton. The substances were originally selected by Munz et al. (2017), who established a list of priority substances based on a large survey in 24 Swiss streams receiving wastewater effluents. The selection criteria included detection frequency, concentrations in municipal wastewater, toxicity, analytical limitations and substance groups. Passive samplers from the channels were prepared and extracted according to Moschet et al. (2015) with few modifications (Carles et al., 2021). Micropollutant concentrations in the passive sampler extracts were then used to derive theoretical concentrations in the water by using the sampling rates (R_S_) established by Moschet et al. (2015). Micropollutants were also analysed in the passive sampler extract used for the community tolerance bioassays – see Tlili et al. (2017) for the detailed description of the extraction method. Concentrations of micropollutants that accumulated in periphyton were measured after extraction by a Quick, Easy, Cheap, Effective, Rugged and Safe (QuEChERS) method as described by Munz et al. (2018) with some modifications (Carles et al., 2021). Micropollutant analysis in all samples was performed by HPLC-MS/MS as described by Carles et al. (2021). The limit of quantification (LOQ with matrix factor correction) and relative recovery for each substance in each type of sample are reported in Table S1.

### 2.5. Biological analyses

#### 2.5.1. Periphyton characterization

Total biomass was determined as ash free dry weight (AFDW) as described in Tlili et al. (2008). Chlorophyll-a content was used as a proxy for algal biomass (Sartory and Grobbelaar, 1984). Bacterial biomass was estimated with flow cytometry according to Frossard et al. (2012) with few modifications (Carles et al., 2021). Freeze-dried periphyton samples were analysed for total carbon and total nitrogen by using an elemental analyser (HEKAtech Euro Elemental Analyzer; HEKAtech GmbH, Wegberg, Germany). Total phosphorus in freeze-dried periphyton was determined by inductively coupled plasma mass spectrometry (8900 Triple Quadrupole ICP-MS, Agilent) after an additional digestion step. This consisted of mixing 5 mg of each periphyton sample with 3 mL of 65 % HNO_3_ and 1 mL of H_2_O_2_ and heating at 250 °C (pressure 120 bars) for 5 min in an ultraCLAVE 4 (Milestone GmbH, Germany).

Photosynthetic efficiency was assessed by using an Imaging-PAM (pulse amplitude-modulated) fluorimeter (Heinz Walz GmbH, Germany). Primary and secondary productions were measured via ^14^C-carbonate incorporation rate and ^14^C-leucine incorporation into protein as described in Dorigo and Leboulanger (2001) and Buesing and Gessner (2003), respectively, with few modifications (Carles et al., 2021). Basal heterotrophic respiration was measured by using the MicroResp™ technique, following the procedure described in Tlili et al. (2011). The same method was also used to determine the community-level physiological profile (CLPP) of the periphyton suspensions from the various treatments. This was done by measuring the mineralization potential of 14 different carbon sources corresponding to 3 amino acids (glycine, L-cysteine and L-serine), 8 carbohydrates (D(+)-glucose, D(+)-xylose, L-arabinose, D(-)-fructose, D(+)-galactose, D(+)-mannose, D-sorbitol and sucrose), 2 carboxylic acids (citric acid and D(+)-galacturonic acid, monohydrate) and 1 organosulfonic acid (MOPS).

#### 2.5.2. Community tolerance bioassays

Tolerance to micropollutants from the wastewater was determined in periphyton from each treatment via short-term exposure assays with serial dilutions of extracts from the passive samplers that had been deployed in wastewater as described in Carles et al. (2021) with few modifications. Briefly, a logarithmic series of six dilutions of the pure micropollutant extract was freshly prepared from the stock solution in Evian mineral water, using a dilution factor of 3.16. This resulted in the following relative dilution factor (RDF, arbitrary unit) of the micropollutant extract: 1000 (pure extract), 313, 98, 31, 10 and 3. Fifty mL periphyton suspensions were prepared from the stock periphyton suspension by adjusting the optical density at 685 nm to 0.4. An aliquot of 4.5 mL of each suspension was then exposed in 20 mL glass vials (Econo glass vials with Foil-Lined Urea Screw Cap, PerkinElmer, Switzerland) to 0.5 mL of each of the six dilutions of the extract for 4 h. In addition, two controls were prepared: One consisted of the periphyton suspension and 0.5 mL of mineral water (chemical-free control) and the second of periphyton and 0.5 mL of 37% formaldehyde (i.e., formaldehyde control), the latter being used to determine the background activity. Subsamples from each vial were taken for photosynthetic efficiency, primary production and bacterial secondary production measurements, respectively, as described above. The same procedure was applied for substrate-induced respiration measurements by using the MicroResp™ technique. An aliquot of 0.5 mL of each suspension was exposed in 96-deepwell microplates to 50 μL of each of the six dilutions of the extract for 14 h, in addition to the two controls describe above.

#### 2.5.3. Next generation sequencing for prokaryotic and eukaryotic community compositions

Total genomic DNA extraction, library construction and sequencing of the 16S rRNA gene for prokaryotes and the 18S rRNA gene for eukaryotes were carried out as described by Carles et al. (2021) with few modifications. Briefly, total genomic DNA was extracted from an aliquot of 2 mL from each biofilm suspension, stream water, and unfiltered or ultra-filtered wastewater (membrane disc filters) by using the DNeasy PowerBiofilm Kit (QIAGEN) following the manufacturer’s instructions. The library construction consisted in a two-step PCR process. The first PCR amplified the V3-V4 region of the 16S rRNA gene for prokaryotes and the V4-V5 region of the 18S rRNA gene for eukaryotes using the primer sets described by Herlemann et al. (2011) and Hugerth et al. (2014), respectively (Table S2). The second PCR was carried out to add multiplexing indices and Illumina sequencing adapters. The libraries were then normalized, pooled and sequenced (paired end 2 × 300 nt, Illumina MiSeq) following the manufacture’s run protocols (Illumina, Inc.). All raw sequences are available at the National Center for Biotechnology Information (NCBI) under the SRA accession ID PRJNA755072.

Sequencing data processing, Amplicon Sequence Variants (ASVs) binning and taxonomic assignment were done according to Carles et al. (2021) with few modifications. Briefly, the reads were checked for quality and end-trimmed by using FastQC v0.11.2 (Andrews, 2010) and seqtk (https://github.com/lh3/seqtk), respectively, and then merged using FLASH v1.2.11 (Magoč and Salzberg, 2011). The primers were trimmed by using cutadapt v1.12 (Martin, 2011). Quality filtering was performed with PRINSEQ-lite v0.20.4 with a subsequent size and GC selection step (Schmieder and Edwards, 2011). The reads were processed with an ASV analysis (Callahan et al., 2017). The sample reads were first denoised into ASVs with UNOISE3 in the USEARCH software v.11.0.667. Final predicted taxonomic assignments were performed with the SILVA v128 (16S rRNA) and the PR2 v4.14.0 (18S rRNA) sequence databases by using SINTAX in the USEARCH software v.11.0.667 (Edgar, 2016). The total number of reads obtained at each bioinformatics step is reported in Table S3.

### 2.6. Data analyses

Significant differences among the treatments for the periphyton descriptors (i.e., AFDW, chlorophyll-a content, bacterial biomass, photosynthetic efficiency, primary production, secondary production, SIR, basal respiration, C:N:P molar ratios and taxonomic abundance) were assessed using one-way ANOVA followed by separate post hoc comparisons (Tukey’s test, α =□0.05). The tested factor was the treatment (five modalities: 0% WW, 30 % WW, 30 % UF, 80 % WW and 80 % UF). Normality and homogeneity of variance were checked prior to ANOVA analysis (Kolmogorov-Smirnov’s and Levene’s tests, respectively, α = 0.05). Data that were not normally distributed were transformed using logarithmic or Box-Cox functions. Statistical analyses were carried out in R 4.0.4 by using RStudio (version 1.4.1717). The effective concentrations leading to a 50% decrease of the activity (EC_50_) in the community tolerance bioassays were derived from concentration-activity relationships as described in Carles et al. (2021). Briefly, concentration-activity relationships of the four biological replicates (N = 4 channels) of each treatment were plotted as a function of decreasing passive sampler extract dilutions. For each treatment, the data corresponding to four biological replicates and 6 passive sampler dilutions (n = 24) were then fitted with the DoseResp function of the OriginPro 2016 software (Origin Lab Corporation, USA). An arbitrary value of RDF = 1000 was set for the pure passive sampler extract.

Sequencing data analyses were performed with the R package *phyloseq* version 1.34.0 (McMurdie and Holmes, 2013). A total of 499,968 and 72,072 reads was obtained after rarefaction for 16S and 18S rRNA datasets, respectively. Alpha diversity (i.e., richness and evenness of a given community) was evaluated for each periphyton and water sample via Chao1 species richness and Shannon diversity index with the package *phyloseq*. The analysis of beta diversity (i.e., measuring the structural differences among several communities) was based on Bray-Curtis distances, which use the relative abundance of taxa. Permutational Multivariate Analysis of Variance (PERMANOVA) tests were carried out on the Bray-Curtis distances matrix of prokaryotic and eukaryotic communities using the R package *vegan*. After testing homogeneity in prokaryotic and eukaryotic datasets dispersion, the *adonis* function was used to test the null hypothesis to see if experimental treatments shared similar centroids. Additional pairwise comparisons were carried out by using the *pairwise.adonis2* function (Martinez Arbizu, 2020). Graphical representations were generated with the R package *ggplot2* version 3.3.5.

The relative contributions of *source* communities (i.e. stream water, wastewater and UF wastewater) were determined for *sink* communities (i.e. periphyton communities) according to the mixture of stream water and wastewater in the channels by using the fast expectation-maximization for microbial source tracking (FEAST) package in R (Shenhav et al., 2019) for both prokaryotes and eukaryotes. The analysis was repeated five times (1000 iterations each) to reduce the effect of false predictions, with 12 replicates (sampling times) for each water *source* and four channel replicates for each *sink* periphyton community. The FEAST analysis also reports on the potential proportion in the *sink* community attributed to other origins, collectively referred as the unknown source.

In order to identify taxa (ASVs) in periphyton responding positively or negatively to wastewater microorganisms, microbiome differential abundance testing was carried out for prokaryotes and eukaryotes by using the R package *DESeq2* version 1.30.1 (Love et al., 2014). This led to the selection of taxa that were assigned to three different groups for prokaryotes and eukaryotes (Figure S1). *Group Positive direct* contains taxa that were positively impacted in periphyton by wastewater microorganisms and have been removed from the wastewater by ultra-filtration. Hence, taxa belonging to *Group Positive direct* have a higher abundance in periphyton 30 and 80 % WW than in periphyton 0 % WW and in periphyton 30 and 80 % UF, respectively, as well as a higher abundance in wastewater than in UF wastewater. *Group Positive indirect* corresponds to taxa that were positively impacted by wastewater microorganisms and did not originate from the WWTP. Therefore, taxa of *Group Positive indirect* have a higher abundance in periphyton 30 and 80 % WW than in periphyton 0 % WW and in periphyton 30 and 80 % UF, respectively, but they do not have a higher abundance in wastewater than in UF wastewater. *Group Negative* contains taxa that were negatively impacted by wastewater microorganisms. These taxa have a lower abundance in periphyton 30 and 80 % WW than in periphyton 0 % WW and in periphyton 30 and 80 % UF, respectively. The differential abundance was tested using the nonrarefied ASV counts. Wald test and adjusted p-values were used to determine if each calculated log2 fold-change differed significantly from zero. In our study, we considered differentially abundant taxa with a log2 fold-change ≥ |2| and Benjamini-Hochberg adjusted p-values < 0.05. The control (0 % WW), 30 and 80 % UF periphyton, as well as the UF wastewater were used as reference. Correlation among the prokaryotic and eukaryotic taxa selected by the microbial differential abundance testing was subsequently assessed based on the relative abundance (variance stabilizing transformation – vst-counts) of each taxon in periphyton from all treatments (N = 20). The correlation matrix was visualized with a heatmap displaying the Pearson (*r*) correlation coefficient (P < 0.05) with the R package *pheatmap* 1.0.12 (Kolde, 2015).

## 3. Results and discussion

### 3.1. Micropollutants in water and in periphyton

As we expected, the measured concentration of each quantifiable substance in the water in the channels increased with wastewater proportion, indicating that wastewater constituted the primary source of micropollutants in the channels (Figure 1A and Table S4). Our results also show that these concentrations were similar between the unfiltered and ultra-filtered wastewater for both 30 and 80 % wastewater proportions (Figure S2). This finding confirms that the ultra-filtration, which was primarily intended to remove microorganisms from the effluent, did not modify the micropollutant composition and concentrations in the wastewater.

**Figure 1.**
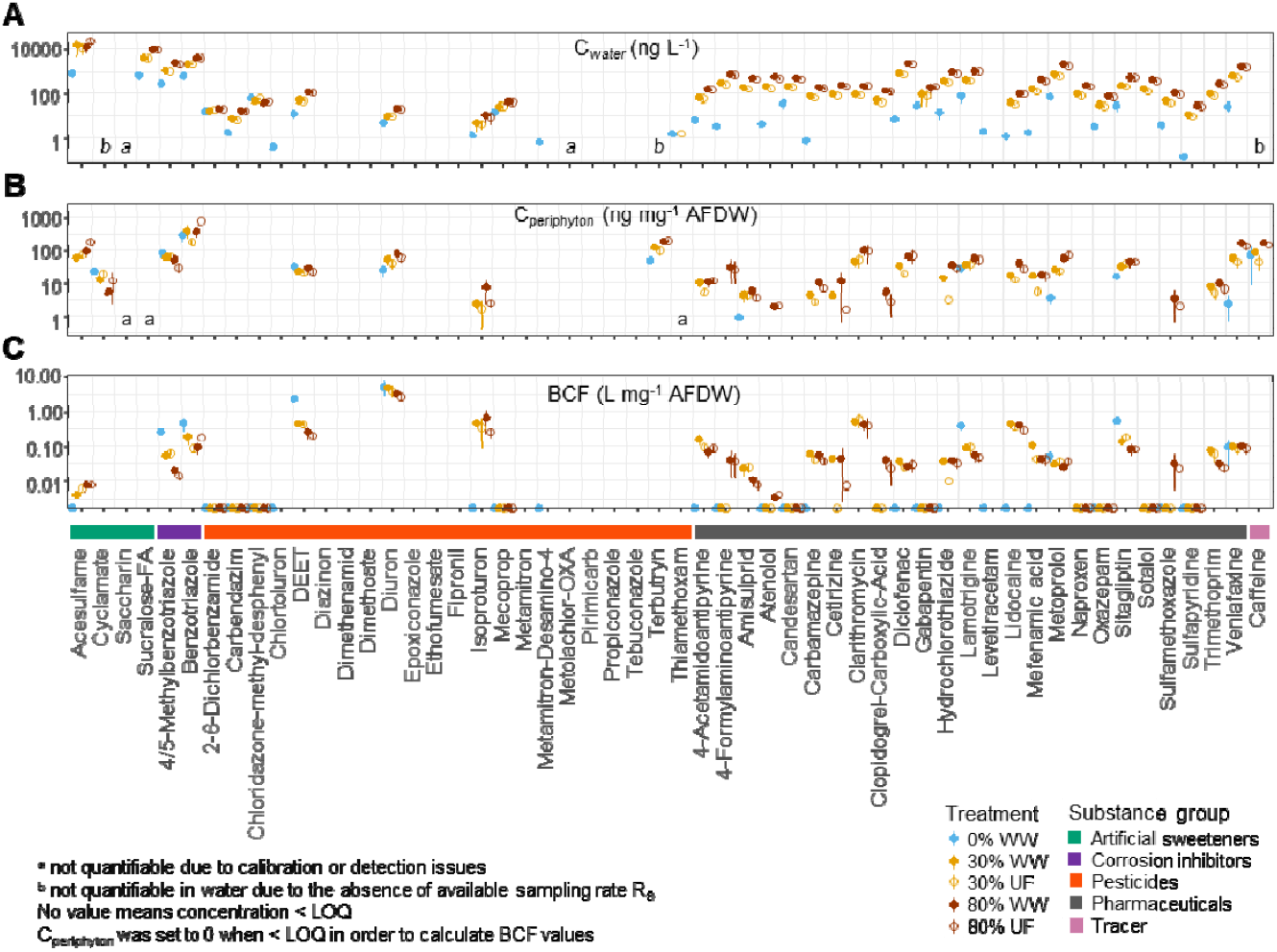
Micropollutant concentration in the channel water and in periphyton. Fifty-four substances were included in the targeted mass spectrometry analysis, including 4 artificial sweeteners, 2 corrosion inhibitors, 22 pesticides, 25 pharmaceuticals and one tracer (caffeine). **A**. Micropollutant concentration in water samples (Cwater) in ng L^-1^. **B**. Micropollutant concentration in periphyton samples (C_periphyton_) in ng mg^-1^ of periphyton ash-free dry weight (AFDW). **C**. Bioconcentration factor (BCF), calculated as the ratio between the concentration in periphyton (in ng mg^-1^ AFDW) and the average concentration in water (in ng L^-1^) for each substance and treatment. C_periphyton_, C_water_ and BCF were reported for each substance and each treatment. The treatments correspond to periphyton grown in the presence of 0% (control), 30% and 80% of unfiltered (WW) and ultra-filtered (UF) wastewater. Values are mean ± SD from 4 channel replicates (C_periphyton_) and 4 passive samplers (C_water_).

Because it is important to link effects to the actual exposure concentrations, we also quantified the micropollutants that accumulated in periphyton (Figure 1B and Table S5). We found micropollutants to accumulate in periphyton with a positive correlation between micropollutant concentrations in periphyton and in the water (Figure S3A), indicating that periphyton was able to accumulate micropollutants from the water phase. However, accumulation varied depending on the measured substance as illustrated by bioconcentration factors (BCFs), which were derived by dividing the concentration in the periphyton by the concentration in the water. The highest BCF values were found for the pesticides diuron, DEET and isoproturon, as well as for some pharmaceuticals such as clarithromycin and lidocaine (Figure 1C and Table S6), a pattern that we also observed in a previous study using the same experimental system and treated wastewater (Carles et al., 2021). Differences in bioaccumulation could be explained by differences in the physicochemical properties of the compounds. This was reflected in the positive correlation that we found for the BCFs and log-transformed octanol/water partition coefficients (LogKow) (Figure S3B). The comparatively high bioaccumulation of PSII inhibitors, such as the herbicides diuron and isoproturon, might be also explained by the presence of specific molecular binding sites, such as the protein D1 in the photosynthetic apparatus, within phototrophic organisms that dominate periphyton (Allen et al., 1983; Morin et al., 2018; Tlili et al., 2011b). On the contrary, the relatively low BCF values for other substances, such as the artificial sweetener acesulfame, may be explained by their high biotransformation rates within microbial cells in periphyton (Desiante et al., 2022, 2021).

### 3.2. Periphyton characterization

We found no significant effect of wastewater, whether filtered or not, on most of the traditional periphyton descriptors, namely biomass, photosynthetic efficiency or primary and secondary production (Table 1), as also reported previously (Carles et al., 2021; Lebkuecher et al., 2018; Pereda et al., 2019; Tlili et al., 2017). On the other hand, basal respiration in the periphyton exposed to 80% of unfiltered and ultra-filtered wastewater significantly increased, indicating a potential shift towards heterotrophy. This conclusion is further supported by the clear effect of wastewater on nutrient stoichiometry. Regardless of the treatment, periphyton exposed to wastewater was characterized by lower C:N and C:P molar ratios than the control, with values closer to those described for bacteria than for algae (Table 1). For instance, heterotrophic bacterial cells were described to have a C:P ratio of about 45 (Goldman et al., 1987) while algal C:P ratio is more around 106 (Redfield et al., 1963). Moreover, bacteria have been shown to possess a higher ability to assimilate inorganic phosphorus than algae (Currie and Kalff, 1984). The shift towards heterotrophy could therefore also be explained by changes in the nutrient composition of the extracellular matrix of periphyton, which is in turn due to the increase of available nutrients in the water phase due to the presence of wastewater. This was confirmed by the water analyses in the channels, which shoed increased concentrations of *ortho*-phosphate, nitrate and dissolved organic carbon in both 80% unfiltered and ultra-filtered wastewater treatments compared to those in the control treatment (i.e. 0% wastewater) (Desiante et al., 2022).

**Table 1.** Descriptors of periphyton from the five experimental treatments. Data are means ± standard deviation from four replicate channels per treatment (N = 4). Significant differences between treatments are indicated by lower case letters (a < b < c, Tukey’s test, P < 0.05). The treatments correspond to periphyton grown in the presence of 0% (control), 30% and 80% wastewater (WW) and ultra-filtered wastewater (UF). Significant difference between wastewater treatments and the control (0% wastewater) is marked in bold.

|  | 0% WW | 30% WW | 30% UF | 80% WW | 80% UF |
| --- | --- | --- | --- | --- | --- |
| Biomass |  |  |  |  |  |
| Ash-free dry weight (mg cm<sup>-2</sup>) | 0.48 ± 0.06 (ab) | 0.54 ± 0.05 (b) | 0.41 ± 0.05 (ab) | 0.41 ± 0.06 (ab) | 0.37 ± 0.03 (b) |
| Chlorophyll-a (mg g<sup>-1</sup> AFDW) | 14.4 ± 1.8 (a) | 15.9 ± 1.5 (a) | 16.6 ± 2.7 (a) | 16 ± 3.5 (a) | 19.6 ± 4.5 (a) |
| Bacterial biomass (μg C g<sup>-1</sup> AFDW) | 1.5 ± 0.6 (a) | 1.5 ± 0.3 (a) | 2.3 ± 0.5 (a) | 2 ± 0.3 (a) | 2.2 ± 0.4 (a) |
| Nutrient ratio |  |  |  |  |  |
| Carbon:Nitrogen molar ratio | 9.5 ± 1.9 (b) | **6.8 ± 0.4 (a)** | **6.3 ± 0.4 (a)** | **5.9 ± 0.4 (a)** | **5.5 ± 0.4 (a)** |
| Carbone:Phosphorus molar ratio | 113.3 ± 12.9 (b) | **61 ± 17.6 (a)** | **57.7 ± 8.3 (a)** | **60.5 ± 23.3 (a)** | **64.1 ± 10.5 (a)** |
| Nitrogen:Phosphorus molar ratio | 12.5 ± 3.5 (a) | 9 ± 2.6 (a) | 9.1 ± 1.3 (a) | 10.1 ± 3.2 (a) | 11.6 ± 1.7 (a) |
| Functional endpoints |  |  |  |  |  |
| Photosynthetic efficiency (Quantum yield □') | 0.36 ± 0.03 (a) | 0.35 ± 0.11 (a) | 0.38 ± 0.02 (a) | 0.42 ± 0.01 (a) | 0.29 ± 0.16 (a) |
| Primary production (μg C g<sup>-1</sup> AFDW day<sup>-1</sup>) | 201.7 ± 49.8 (a) | 210.7 ± 80.9 (a) | 208.5 ± 81.4 (a) | 320.8 ± 49.5 (a) | 144.8 ± 125.7 (a) |
| Secondary production (μg C g<sup>-1</sup> AFDW day<sup>-1</sup>) | 7.9 ± 6.8 (ab) | 4.2 ± 5.3 (ab) | 13.9 ± 6.8 (b) | 3.5 ± 4.1 (ab) | 1.5 ± 0.7 (a) |
| Basal respiration (g CO2 g<sup>-1</sup> AFDW day<sup>-1</sup>) | 3.8 ± 0.3 (a) | 3.9 ± 0.5 (ab) | 4.1 ± 0.6 (abc) | **5.3 ± 0.8 (c)** | **5.2 ± 0.6 (bc)** |

Impacts of wastewater on the heterotrophic component of periphyton were also reflected by the changes in the community-level physiological profiles (CLPPs) that we established based on the capability of heterotrophs to mineralize various carbon sources for respiration (Figure 2). Indeed, the respiration profiles of periphyton exposed to unfiltered and ultra-filtered wastewater differed from each other and from the control communities along both PCA 1 and PCA 2 axes. The PCA 1 is clearly related to the wastewater proportion in the channels and shows a positive correlation between wastewater increase and a higher mineralisation potential of all tested carbon substrates. Such results underline again the fact that wastewater exposure favoured the increase of heterotrophic activities in periphyton communities. Interestingly, the dissimilarity along PCA 2 between the communities exposed to the ultra-filtered wastewater and the control was less pronounced than for periphyton exposed to the unfiltered wastewater. This suggests that microorganisms in the wastewater have contributed to some extent to the observed changes. It is plausible that such microorganisms have the capacity to degrade all types of organic substrates in the highly enriched activated sludge compartment of the WWTP, before being released with the effluent and colonizing periphyton.

**Figure 2.**
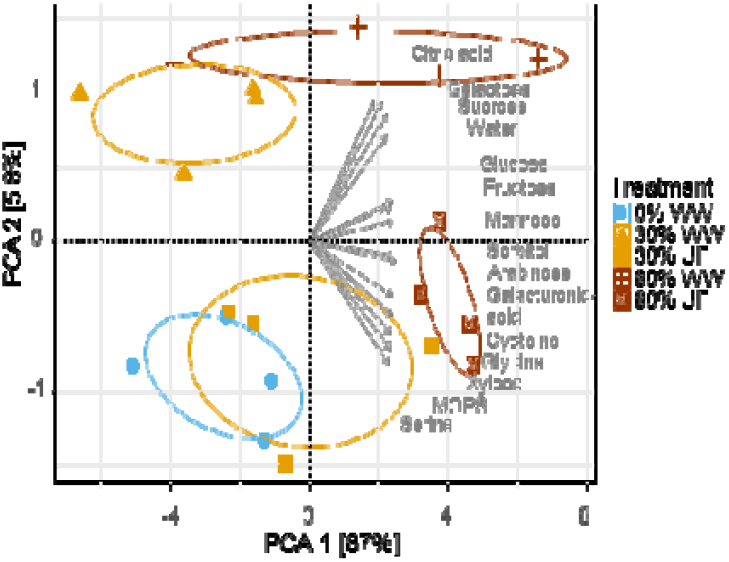
Principal component analysis (PCA) of the community-level physiological profiles (CLPP) of heterotrophs in periphyton. The profiles were established based on the capacity of heterotrophs to mineralize various carbon sources for respiration, including three amino acids, eight carbohydrates, two carboxylic acids and one organosulfonic acid, as well as a control without an additional carbon source (i.e. measurement of basal respiration). The variables were normalized by the function PCA() of the R package *FactoMineR*. The treatments correspond to periphyton grown in the presence of 0% (control), 30% and 80% of unfiltered (WW) and ultra-filtered wastewater (UF) wastewater. The 95 % confidence ellipse was added for each treatment. Non-overlapping ellipses indicate significant differences between treatments.

### 3.3. Tolerance of periphyton to micropollutants

Increased microbial community tolerance in periphyton towards micropollutants from WWTPs has been suggested as an indicator to disentangle the specific effects of micropollutants from those of other stressors (Corcoll et al., 2014; Tlili et al., 2020, 2017). We assessed the tolerance of periphyton to a complex mixture of micropollutants that is representative of the micropollutants found in the wastewater effluent (Figure S4 and Table S7). Irrespective of the treatment, no effect of wastewater was observed on periphyton tolerance based on primary production and respiration, and inconclusive results were obtained when measuring secondary production due to the absence of inhibition and the high variability among replicates (Table 2 and Figure S5). In sharp contrast, calculated EC50 values based on photosynthetic PSII yield measurements were significantly higher for periphyton exposed to 30 and 80 % unfiltered wastewater than for the controls. This reflects an increased tolerance of the phototrophic communities to the micropollutant mixture, as also shown in our previous study (Carles et al., 2021). A potential explanation could be related to the mode of action of PSII inhibitor herbicides, which bind to the QB-binding niche on the D1 protein of the PSII complex, thus blocking electron transport from QA to QB (Jansen et al., 1993). It has been suggested that upon exposure to such herbicides, tolerant phototrophic species can upregulate the expression of the D1 protein, leading to an increased abundance of the QB-PSII inhibitors complex. A community dominated by such tolerant species can therefore cope with higher exposure concentrations than a community dominated by sensitive species, leading to higher EC_50_ values (Tlili et al., 2011b). This is in agreement with the higher BCFs we found for the herbicides PSII inhibitors, such as diuron and isoproturon.

**Table 2.** Tolerance measurements of periphyton from the five experimental treatments. The treatments correspond to periphyton grown in the presence of 0% (control), as well as 30% and 80% of unfiltered (WW) or ultra-filtered wastewater (UF). EC_50_ values were derived from the fitting model applied to the four replicate channels per treatment (N = 4). Higher values for each wastewater treatment than the control means higher tolerance to the micropollutant mixture. The x-axis of the concentration-effect relationships was expressed as a unit-less relative dilution factor (RDF) and therefore the EC_50_ values are also expressed in RDF. Values in parentheses correspond to the 95% confidence interval. Ratios (R) of EC_50_ were calculated for each endpoint by dividing the mean EC_50_ of 30% WW, 30% UF, 80% WW and 80% UF by the corresponding EC_50_ of the control. (R ≤ 1 indicates no induced tolerance; R > 1 induced tolerance); n.d. means not determined due to the absence of inhibition. Significant difference between wastewater treatments and the control (0% wastewater) is marked in bold.

| Endpoint | 0% WW |  | 30% WW |  | 30% UF |  | 80% WW |  | 80% UF |  |
| --- | --- | --- | --- | --- | --- | --- | --- | --- | --- | --- |
|  | EC<sub>50</sub> | R | EC<sub>50</sub> | R | EC<sub>50</sub> | R | EC<sub>50</sub> | R | EC<sub>50</sub> | R |
| Photosynthetic efficiency | 354 (231 - 545) | — | **800 (541 - 1186)** | **2.3** | 348 (213 - 571) | 0.9 | **653 (492 - 869)** | **1.8** | 299 (167 - 536) | 0.8 |
| Primary production | 133 (119 - 149) | — | 139 (108 - 179) | 1 | 151 (116 - 198) | 1.1 | 154 (121 - 196) | 1.2 | 136 (106 - 175) | 1 |
| Secondary production | n.d. | n.d. | n.d. | n.d. | n.d. | n.d. | n.d. | n.d. | n.d. | n.d. |
| Substrate-induced respiration | 54 (32 - 90) | — | 58 (35 - 97) | 1.1 | 63 (37 - 106) | 1.2 | 40 (21 - 76) | 0.7 | 89 (42 - 191) | 1.7 |

Regardless of the underlying mechanisms, an important insight from our study is that following the ultrafiltration of the wastewater, which removed more than 99 % of the microorganisms, no increased tolerance was observed in periphyton (Table 2 and Figure S5). Similarly, Desiante et al. (2022) have reported on a loss of micropollutant biotransformation potential by periphyton, one of the mechanisms potentially explaining tolerance, following the ultrafiltration of the wastewater. Collectively, such findings point towards a major role that microorganisms originating from the WWTPs play in community tolerance to micropollutants. It is conceivable that these microorganisms might have developed a tolerance to micropollutants in the WWTP before being released into the streams. Understanding whether their contribution occurs directly via the colonization of periphyton by micropollutant-tolerant taxa, or indirectly by modifying species interactions within the community, requires a comprehensive characterization of the microbial diversity not only in periphyton but as well in wastewater and stream water.

### 3.4. Microbial diversity and taxonomic abundance in periphyton and water

#### 3.4.1. Relative contribution of stream water and wastewater communities to the periphyton community

In order to estimate the relative contribution of the *source* communities (stream water and wastewater) to each *sink* community (periphyton), we used the microbial source tracking tool FEAST (Shenhav et al., 2019). Our results clearly indicated that wastewater contributed largely to periphyton communities, with a higher proportion for prokaryotes than for eukaryotes (Figure 3). For instance, the relative proportion of wastewater in periphyton exposed to 80 % wastewater reached up to 79 % and 38 % for prokaryotes and eukaryotes, respectively. This is in line with the shift towards heterotrophy we observed in periphyton exposed to wastewater, since prokaryotes in periphyton correspond mainly to heterotrophic bacteria. Another line of evidence comes from the comparison of periphyton exposed to unfiltered and ultra-filtered wastewater: we observed a strong reduction of the relative proportion of wastewater community for periphyton exposed to ultra-filtered wastewater (Figure 3). This led, for instance for eukaryotes, to a proportion of wastewater community ≤ 5 % in periphyton exposed to 30 % and 80 % ultra-filtered wastewater, confirming in turn the capacity of wastewater microorganisms to colonize periphyton.

**Figure 3.**
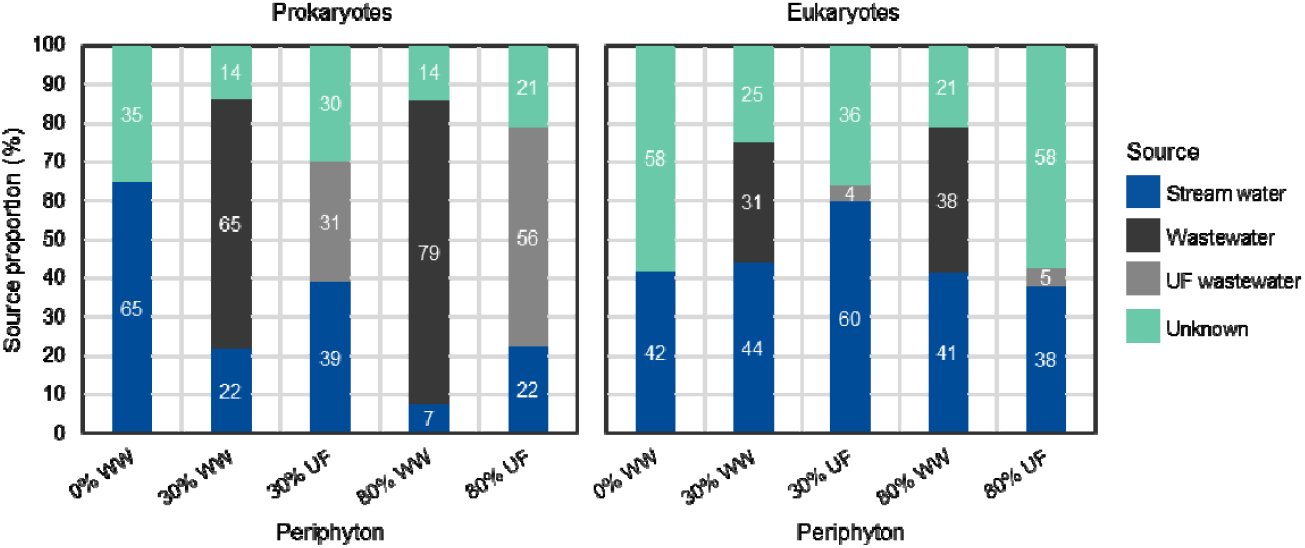
Proportion of water sources in periphyton samples using fast expectation-maximization for microbial source tracking (FEAST) for prokaryotes and eukaryotes. Periphyton was grown in the presence of 0% (control), 30% and 80% of unfiltered (WW) and ultra-filtered (UF) wastewater. The relative contribution of each *source* community (i.e. stream water, unfiltered wastewater and ultra-filtered wastewater) was determined for each *sink* community (i.e. periphyton community) according to the mixture of stream water and wastewater in the channels. FEAST also reports on the potential proportion of the sink attributed to other origins, collectively referred as the unknown source. The analysis was repeated five times (1000 iterations each) with 12 replicates (sampling times) for each water source and four channel replicates for each periphyton community. Data are mean of each source proportion (five independent repetitions).

The proportion of each *sink* community that did not match the signature of the *sources* included in our analysis was assigned to *unknown sources*. Such analysis is used to identify potential contamination of the *sinks* by other unidentified microbial *sources* (Liang et al., 2021; Shenhav et al., 2019). In our study, ultra-filtration did not only lead to an increase of the relative proportion of stream communities in periphyton, but also to an increased proportion of these *unknown sources*. The relatively high proportion of the *unknown sources* may be explained by the presence of microorganisms in the Maiandros channel system, and more specifically for the UF treatment, by the colonization of the backside of the membranes by microorganisms forming distinct communities. The colonization dynamics *per se*, as well as species interactions, in periphyton communities could also lead to different community assemblages compared to the surrounding water column (Peng et al., 2018), and thus potentially contribute to the relatively high proportion of the *unknown sources*. This makes periphyton different from free-living microorganisms in the water column in streams, for which several field surveys have shown that downstream bacterial community profiles were a mixture between the upstream and the effluent (Mansfeldt et al., 2020; Pascual-Benito et al., 2020; Price et al., 2018).

#### 3.4.2. Impact of wastewater and wastewater microorganisms on periphyton community structure

We further evaluated the impacts of wastewater on periphyton community structure based on the commonly used descriptors alpha- (i.e. richness and Shannon diversity index of a given community) and beta-diversity (i.e. structural differences among several microbial communities). While alpha-diversity for eukaryotes was not affected by wastewater, both taxonomic richness (Chao1) and Shannon diversity of prokaryotes decreased in periphyton exposed to 80 % wastewater compared to the control, with no significant difference between unfiltered and ultra-filtered wastewater treatments (Figure 4A). In contrast, wastewater led to a clear separation in the beta-diversity of both prokaryotic and eukaryotic communities among all treatments (PERMANOVA, *P* = 0.001, Figure 4B). Indeed, periphyton communities exposed to unfiltered and ultra-filtered wastewater were distinct from each other and from the control communities (pairwise PERMANOVA, *P* < 0.05). Several field and mesocosm studies have also reported on the effects of wastewater effluents on the structure of periphyton communities (Carles et al., 2021; Chonova et al., 2019; Lebkuecher et al., 2018; Romero et al., 2019; Tardy et al., 2021), with contrasting results between alpha- and beta-diversity (Carles et al., 2021; Lebkuecher et al., 2018). This may be explained by the various wastewater constituents, such as nutrients, micropollutants and microorganisms. While micropollutants can negatively affect the abundance of certain taxa, nutrients can favour the growth of others. Collectively, this may result in distinct communities with regard to beta-diversity but with similar alpha-diversity indices.

**Figure 4.**
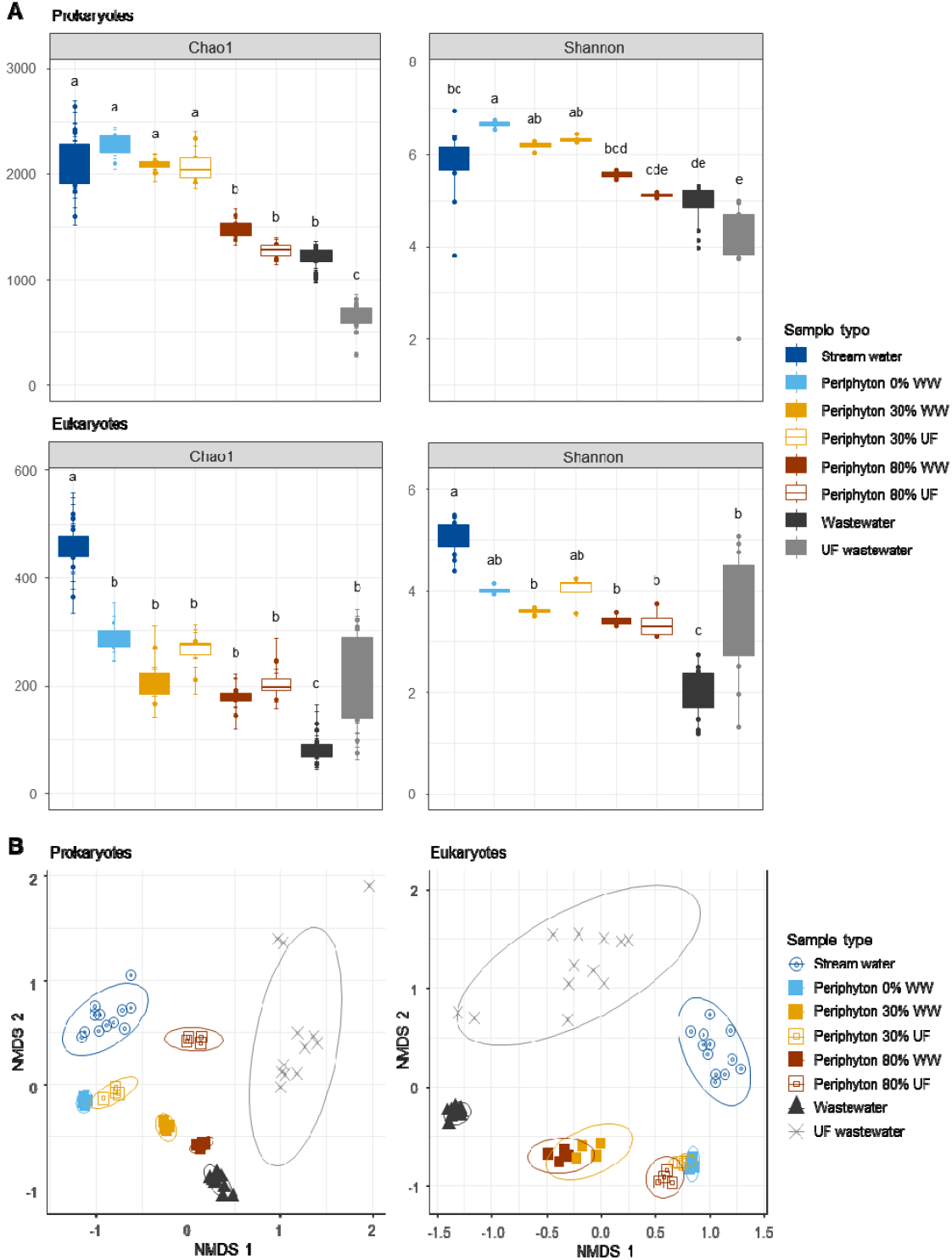
Diversity of prokaryotes and eukaryotes in water and in periphyton. The analyses is based on the next generation sequencing of the 16S rRNA (prokaryotes) and 18S rRNA (eukaryotes) genes from water samples (stream water, unfiltered wastewater (WW) and ultra-filtered-wastewater (UF)) and from periphyton grown in the presence of 0% (control), 30% and 80% unfiltered (WW) and ultra-filtered (UF) wastewater. **A**: Alpha diversity; the values of total ASV richness Chao1 and Shannon’s diversity index H’ are reported as a boxplot for water (N = 12 sampling times) and periphyton samples (N = 4 channel replicates). Significant differences are indicated by lowercase letters, a > b > c > d > e (Tukey’s test, P < 0.05). **B**: Beta diversity; Non-metric Multi-Dimensional Scaling (NMDS) of prokaryotic and eukaryotic communities based on Bray-Curtis distances. The 95 % confidence ellipse was added for each sample type. Non-overlapping ellipses indicate significantly different sample types.

#### 3.4.3. Identification of periphyton taxa impacted by wastewater microorganisms

By comparing periphyton communities exposed to the unfiltered and ultra-filtered wastewater, as well as with the control communities, we were able to examine how and which microorganisms originating from the wastewater potentially affected the final composition of the communities. For this, we applied the microbial differential abundance testing to identify prokaryotic and eukaryotic taxa in periphyton that are positively or negatively impacted by wastewater microorganisms, in terms of relative abundance. This analysis led to the selection of 129 prokaryotic and 20 eukaryotic taxa (Table S8) that were all correlated to each other (Figure S6) and subsequently assigned to three groups.

The first group, named *Group Positive direct*, was composed of 98 prokaryotic and 8 eukaryotic taxa that were removed by the ultra-filtration from the wastewater and directly colonized periphyton exposed to unfiltered wastewater (Figure 5). All these taxa had a higher abundance in periphyton exposed to unfiltered wastewater than in periphyton exposed to the ultra-filtered wastewater and in the control. Taxa composing this group are therefore potentially major players in periphyton metabolic alterations and increased tolerance that we observed in our study. For instance, two prokaryotic phyla (Chlorobi and Firmicutes) that were only found in this group (Figure 5A), have been frequently detected in the outlet and downstream-periphyton of WWTPs (Aubertheau et al., 2017; Ross et al., 2012; Thiel et al., 2019; Ziganshina et al., 2016) as well as in biofilters used to treat urban wastewater (Aguirre-Sierra et al., 2016). Several taxa that belong to the prokaryotic phylum Chloroflexi were also assigned to this group. Chloroflexi is a phylum of filamentous bacteria possessing a wide diversity of metabolisms and ecological roles, but are best known as photoheterotrophs (Overmann, 2008). This phylum was found to be highly abundant in unfiltered wastewater and periphyton exposed to this wastewater, while almost no Chloroflexi was detected in ultra-filtered wastewater and, accordingly, in periphyton exposed to ultra-filtered wastewater (Figure S7A). This suggests that these phototrophic bacteria could have contributed directly or indirectly to the observed increased tolerance of phototrophs in periphyton to micropollutants.

**Figure 5.**
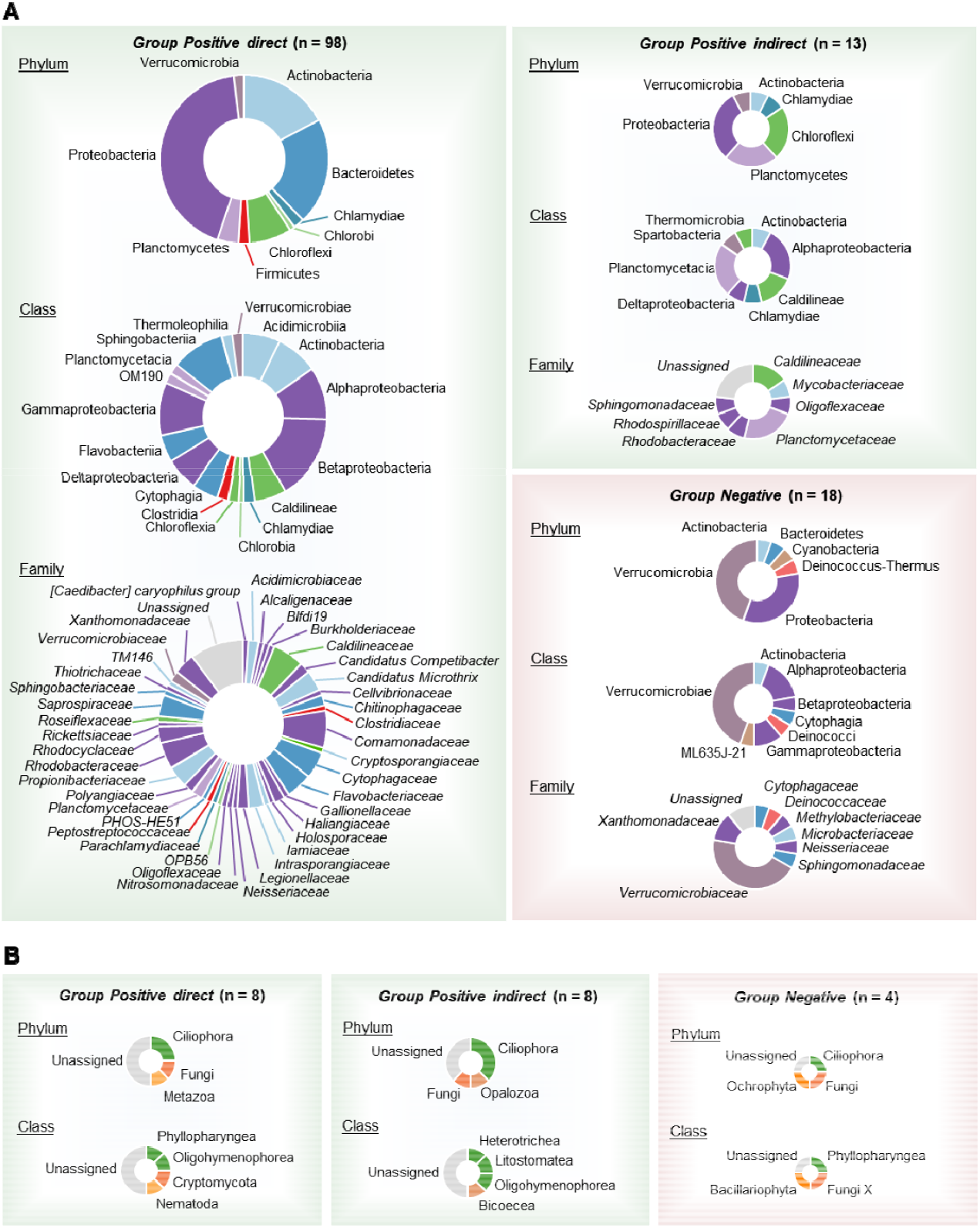
Repartition of the prokaryotic (**A**) and eukaryotic (**B**) taxa selected by the microbial differential abundance testing using DESeq2. The total number of taxa is indicated for each group. The repartition of taxa in each group is described at the phylum, class and family levels for prokaryotes, and at the phylum and class levels for eukaryotes.

The second group, named *Group Positive indirect*, contained 13 prokaryotic and 8 eukaryotic taxa that did not originate from the WWTP but were positively impacted, in terms of relative abundance in periphyton, by the microorganisms from the wastewater (Figure 5). Among the prokaryotic taxa, two families (*Mycobacteriaceae* and *Rhodospirillaceae*) were specific to this group. *Rhodospirillaceae* have been shown to be more abundant in lower and medium order streams (Chen et al., 2018), as it is the case for the stream Chriesbach used in our study. It has also been shown that *Mycobacteriaceae* can thrive in environments that are influenced by anthropogenic activities, such as WWTPs (Amha et al., 2017; Falkinham, 2015; Makovcova et al., 2014). *Group Positive indirect* also contained members of the phylum Chloroflexi, mainly the family *Caldilineaceae* that was also found in the *Group Positive direct* (Figure 5A). *Caldilineaceae* may have contributed to the changes in the respiration profiles of periphyton since it has been shown that this family has different metabolic potentials for substrate utilization compared to other microorganisms involved in the enhanced biological phosphorus removal process of WWTPs (Kindaichi et al., 2013). Microorganisms from the wastewater also had negative impacts on the relative abundance of several taxa in periphyton, which were assigned to the third group, named *Group Negative*. This group was composed by 18 prokaryotic and 4 eukaryotic taxa, among which several were specific to *Group Negative* (Figure 5). For instance, this was the case for several prokaryotic taxa belonging to Cyanobacteria, *Deinococcaceae*, *Methylobacteriaceae* and *Microbacteriaceae* (Figure 5A). Cyanobacteria are known to respond differently (i.e. an increase or a decrease of abundance) to wastewater effluents (Carles et al., 2021; Carles and Artigas, 2020; Corcoll et al., 2014; Mansfeldt et al., 2020; Romero et al., 2019). However, we are not aware of any study that investigated the specific effect of wastewater microorganisms on this phylum. *Deinococcaceae* have already been associated to periphyton growing in reference (unpolluted) sites in a previous study comparing community composition along an urban pollution gradient in a stream (Pineda-Mora et al., 2020), indicating their potential sensitivity to wastewater constituents. The presence of *Methylobacteriaceae* and *Microbacteriaceae* in the *Group Negative* may be explained by their association with microalgae (Levy et al., 2009; Paddock et al., 2020), which seemed to be negatively impacted by wastewater microorganisms in terms of abundance (i.e. Class Chlorophyceae in Figure S7B). Finally, *Group Negative* also contained one eukaryotic taxon affiliated to Bacillariophyta (i.e. diatoms, Figure 5B), which is consistent with the observed negative impact of wastewater microorganisms on diatom abundance (Figure S7B).

These results highlight the need to consider wastewater microbial communities as a stressor *per se*, which can influence the final composition of periphyton communities, when examining the potential impacts of wastewater on stream ecosystems for water quality assessment. This aspect was poorly described so far, as most studies looked either at the overall effluent toxicity (Liao et al., 2019; Nega et al., 2019; Peng et al., 2018; Romero et al., 2019), or at other wastewater constituents, such as nutrients (Lebkuecher et al., 2018) and micropollutants (Carles et al., 2021; Chonova et al., 2019; Tamminen et al., 2022; Tardy et al., 2021; Tlili et al., 2020). Our study also points towards the importance of species interactions within periphyton communities, as well as between prokaryotes and eukaryotes, which occur among co-existing species in periphyton (Gubelit and Grossart, 2020). For instance, studies on diatom-bacterial interactions revealed a high species-specific interdependence of the algal host and bacteria, as each diatom species developed a bacterial community that differed in its composition (Grossart et al., 2005; Koedooder et al., 2019; Stock et al., 2019).

## 4. Conclusions

Overall, our study provides compelling evidence that microorganisms originating from wastewater strongly affected periphyton communities. Specifically, we show that these microorganisms are able to colonize periphyton and modify its community composition, either directly or indirectly via species interactions, contributing to a shift from autotrophy towards heterotrophy. Being at the basis of the food web in streams, such changes in periphyton communities, downstream of WWTPs, bears potential significant environmental costs for higher trophic levels with probable impacts on the overall flow of energy through the food chain in fresh waters. Furthermore, our results also showed that tolerance of periphyton communities to micropollutants was governed by microorganisms released from the WWTP and not by in-stream exposure to the micropollutants. This finding underlines the fact that the Pollution-Induced Community Tolerance (PICT) concept, which stipulates that increased tolerance is directly caused by the exerted selection pressure of micropollutants, should be reconsidered in the context of WWTPs. Instead, selection pressures that occur in highly contaminated compartments such as in the WWTP itself has to be taken into account. Collectively, our findings also have important implications for WWTP management. Microbial communities that are released in the wastewater should be considered as a potential stressor for the receiving streams, similarly to other stressors, such as nutrients, micropollutants or increased temperature. This in turn implies that the measures currently implemented in WWTPs to reduce the load of released microorganisms are not sufficient to completely reduce the ecological hazard they might represent.

## CRediT author statement

**Louis Carles**: conceptualization, data acquisition, formal analysis, writing - original draft, review & editing. **Simon Wullschleger**: data acquisition, formal analysis, review & editing. **Adriano Joss**: funding acquisition, conceptualization, data acquisition, review & editing. **Rik I.L. Eggen**: funding acquisition, conceptualization, review & editing. **Kristin Schirmer**: funding acquisition, conceptualization, review & editing. **Nele Schuwirth**: funding acquisition, conceptualization, review & editing. **Christian Stamm**: funding acquisition, conceptualization, review & editing, project administration. **Ahmed Tlili**: funding acquisition, conceptualization, writing - original draft, review & editing.

## Declaration of interests

The authors declare that they have no known competing financial interests or personal relationships that could have appeared to influence the work reported in this paper.

## Acknowledgements

This work was supported by the project EcoImpact 2.0 funded by Eawag. The authors thank Eawag colleagues Marco Kipf and Richard Fankhauser for their help in setting up the artificial channels, Werner Desiante for his help in maintaining the artificial channels and Bettina Wagner for her help in sampling and periphyton characterization. As well, we would like to thank Aria Minder, Silvia Kobel and Jean-Claude Walser from the Genetic Diversity Centre (GDC, ETH Zürich) for their support with genetic diversity analyses. Data on molecular diversity produced and analysed in this paper were generated in collaboration with the GDC, ETH Zurich.

## Notes

### Competing Interest Statement

The authors have declared no competing interest.

