## Supplemental figures for "Wastewater microorganisms impact microbial diversity and important ecological functions of stream periphyton"

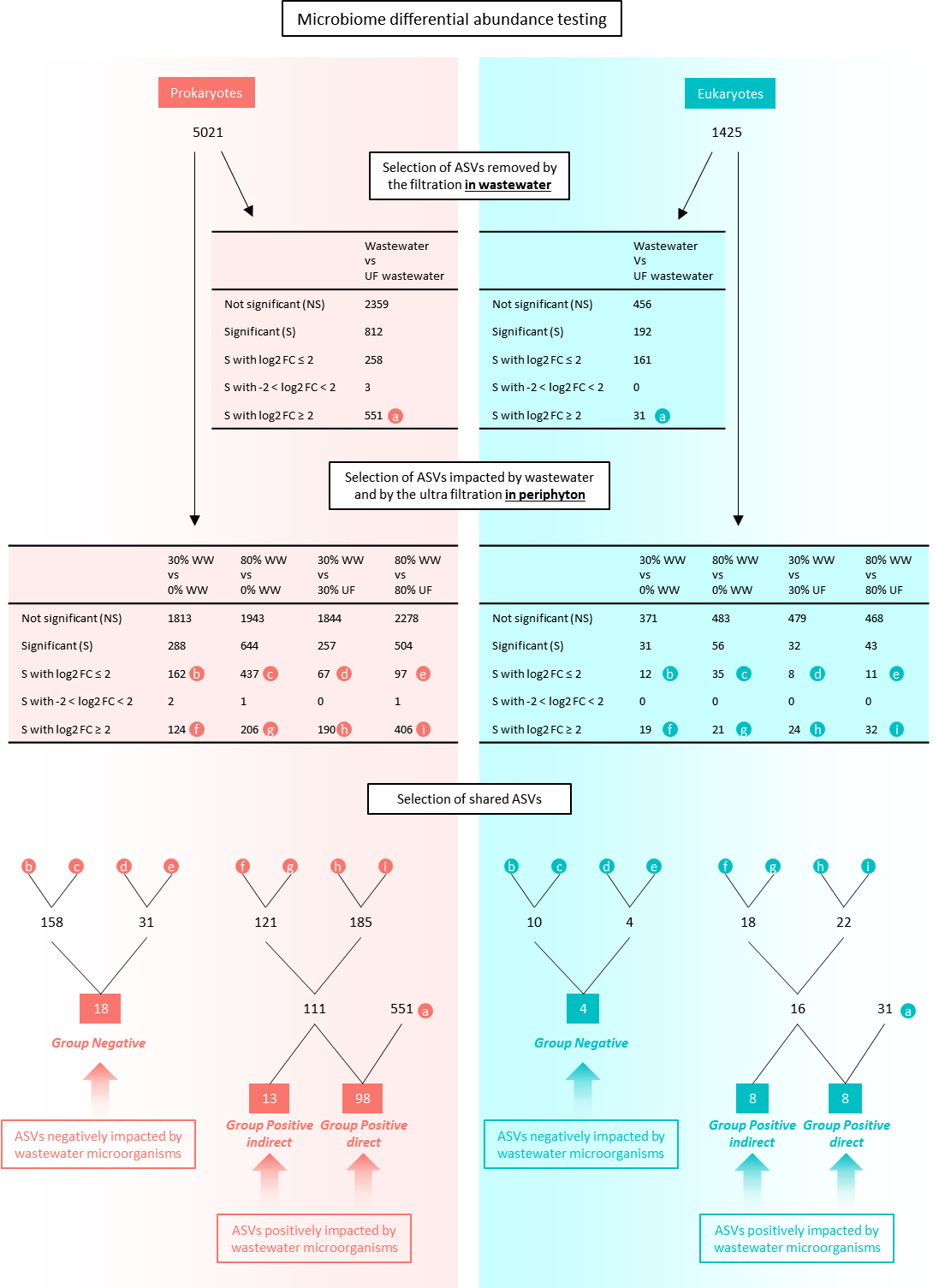

**Figure S1**. Workflow for the microbial differential abundance testing. The same procedure was applied to prokaryotes and eukaryotes datasets. The number of different ASVs is indicated at each step of the analysis. The differential abundance analysis was done with the R package *DESeq2*. The tables show the result of the DESeq2 analyses. Log2 FC: log2 fold-change. The final step of the workflow consisted in the selection of shared taxa between each of the result of the DESeq2 analysis, leading to three groups of taxa for prokaryotes and eukaryotes.

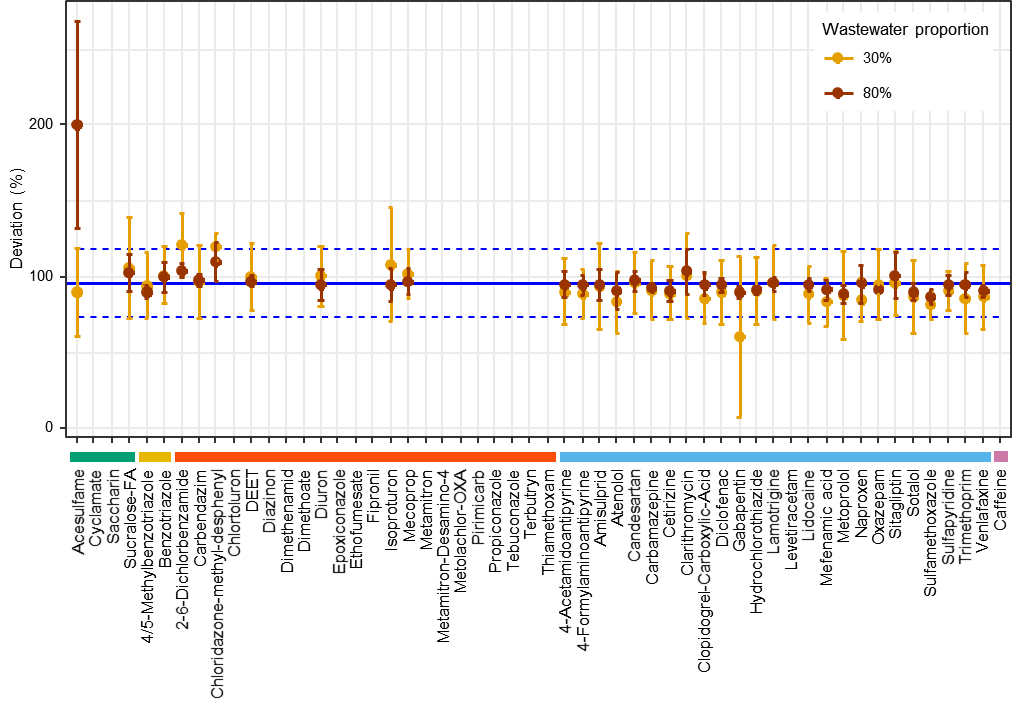

**Figure S2.** Deviation of micropollutant concentration in ultra-filtered wastewater from unfiltered wastewater. The deviation percentage was calculated as the ratio between the concentration in ultra-filtered wastewater and the concentration in unfiltered wastewater for each substance and treatment. Values are mean ± SD from 4 passive samplers for each wastewater proportion (30% and 80%). The blue lines correspond to the average deviation (solid line) ± SD (dashed lines).

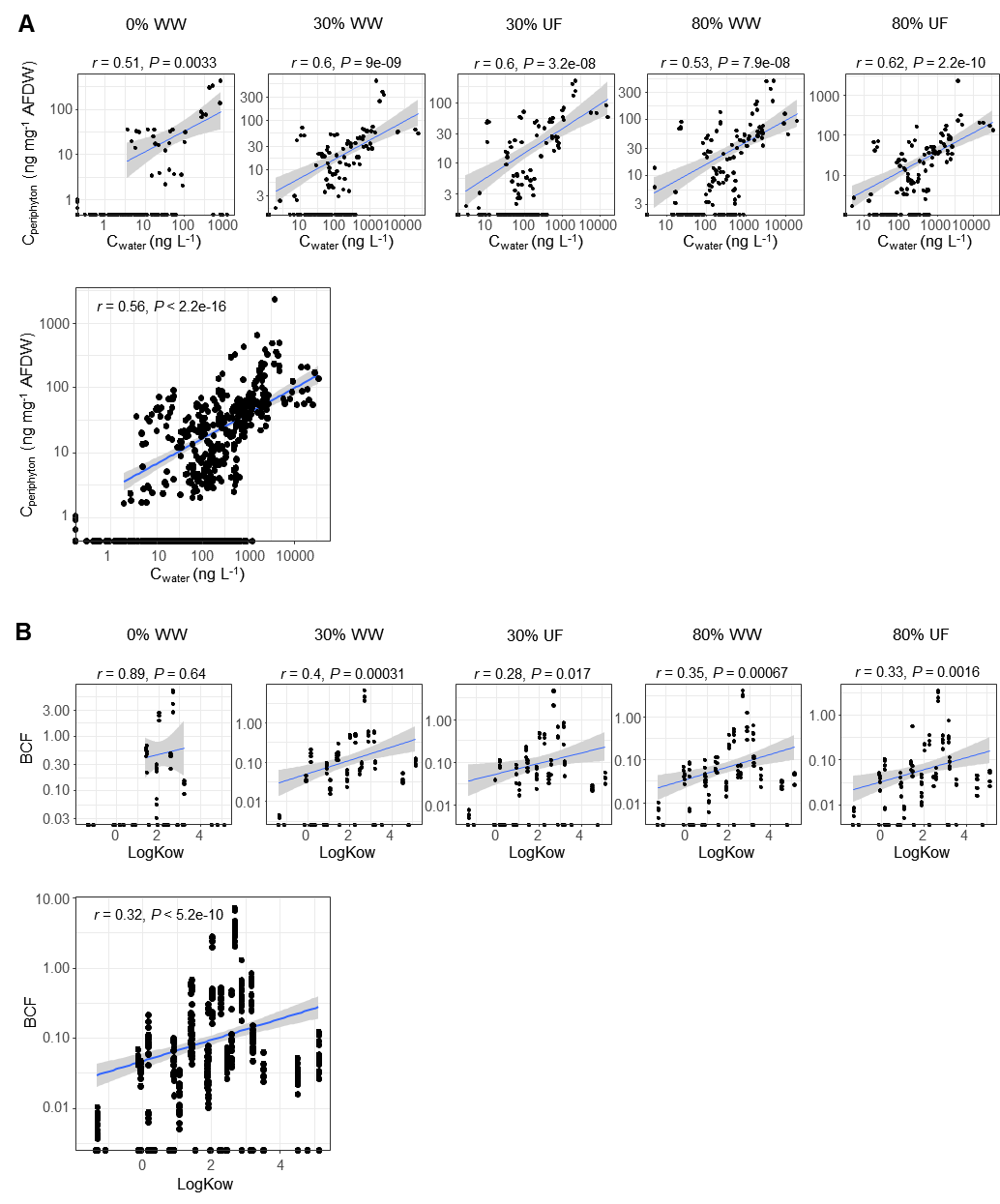

**Figure S3**. Micropollutant concentration in water and in periphyton. Fifty-four substances were included in the targeted mass spectrometry analysis, including 4 artificial sweeteners, 2 corrosion inhibitors, 22 pesticides, 25 pharmaceuticals and one tracer (caffeine). **A**. Correlation between micropollutant concentration in periphyton samples (C_periphyton_) and in water samples (C_water_) for each treatment and for all treatments together. **B**. Correlation between the bioconcentration factor (BCF) and log-transformed octanol/water partition coefficient (LogKow) for each treatment and for all treatments together. The treatments correspond to periphyton grown in the presence of 0% (control), as well as 30% and 80% of unfiltered (WW) and ultra-filtered (UF) wastewater. The blue line corresponds to the linear fit of the log-transformed data. The gray band corresponds to the 95% confidence interval. Values displayed above each plot correspond to the Pearson (*r*) correlation coefficient and the associated p-value (*P*).

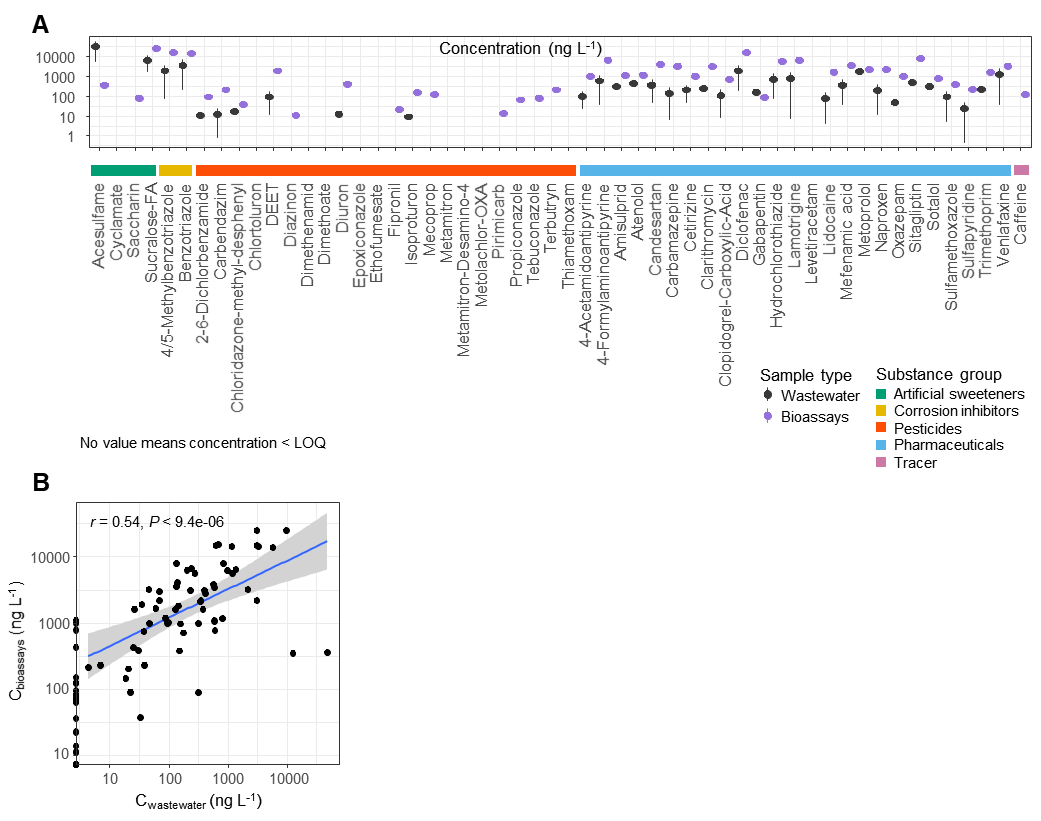

**Figure S4.** Micropollutant concentration in wastewater samples (Wastewater) and in the extract used for the community tolerance bioassays (bioassays) in ng L^-1^. Fifty-four substances were included in the targeted mass spectrometry analysis, including 4 artificial sweeteners, 2 corrosion inhibitors, 22 pesticides, 25 pharmaceuticals and one tracer (caffeine). **A**. Micropollutant concentrations. Bioassays: an arbitrary value of relative dilution factor (RDF) = 1000 was set for the pure passive sampler extract. The results reported here correspond to an RDF of 3. Values are mean ± SD from 2 passive samplers (Wastewater) and 2 dilution replicates (bioassays). **B**. Correlation between the concentrations of micropollutants in wastewater samples (C_Wastewater_) and in the extract used for the bioassays (C_bioassays_). The blue line corresponds to the linear fit of the log-transformed data. The gray band corresponds to the 95% confidence interval. Values displayed above the plot correspond to the Pearson (*r*) correlation coefficient and the associated p-value (*P*).

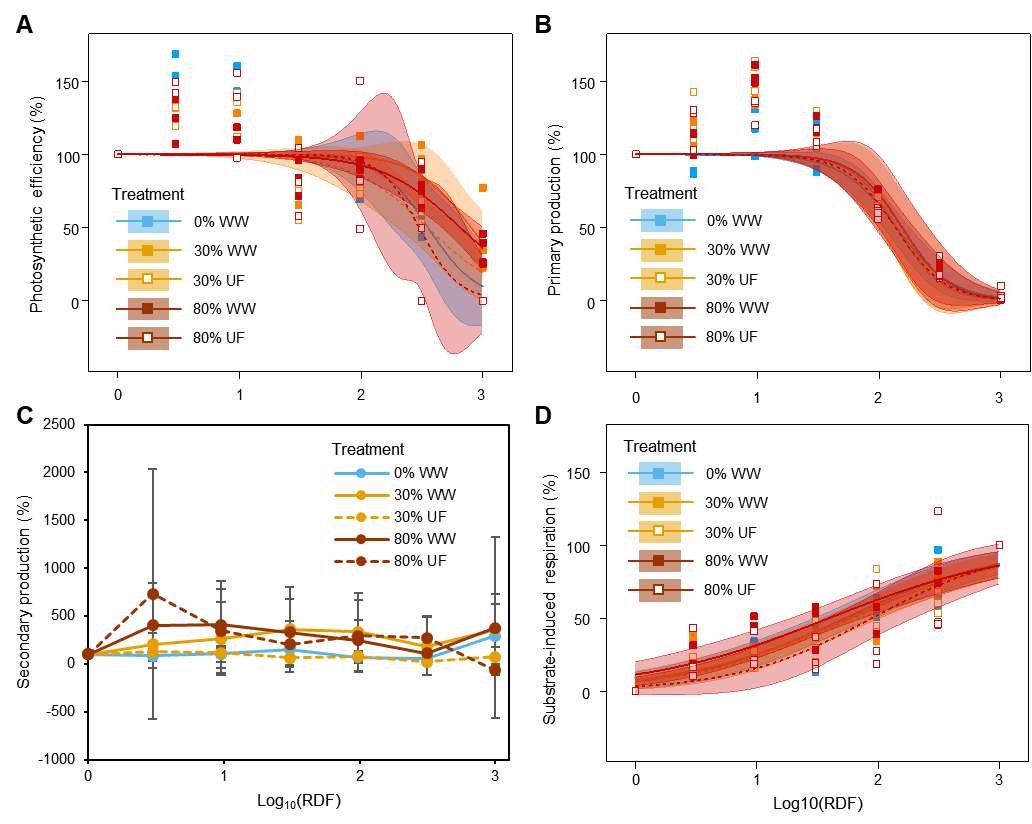

**Figure S5**. Concentration-effect curves based on photosynthetic efficiency (**A**), primary production (**B**), secondary production (**C**) and substrate-induced respiration (**D**) after exposure of periphyton to a serial dilution of the passive sampler extract. The x-axis is expressed in log_10_(relative dilution factor): the maximum value of the RDF, which corresponds to the pure extract, was arbitrary set to 1000 (arbitrary unit). The treatments correspond to periphyton grown in the presence of 0% (control), as well as 30% and 80% of unfiltered (WW) and ultra-filtered (UF) wastewater. **A**, **B** and **D**: four independent replicates are represented together with fitting lines and 95% confidence bands corresponding to the dose-response function described in the Material and Methods section. **C**: data are mean ± SD (n = 4).

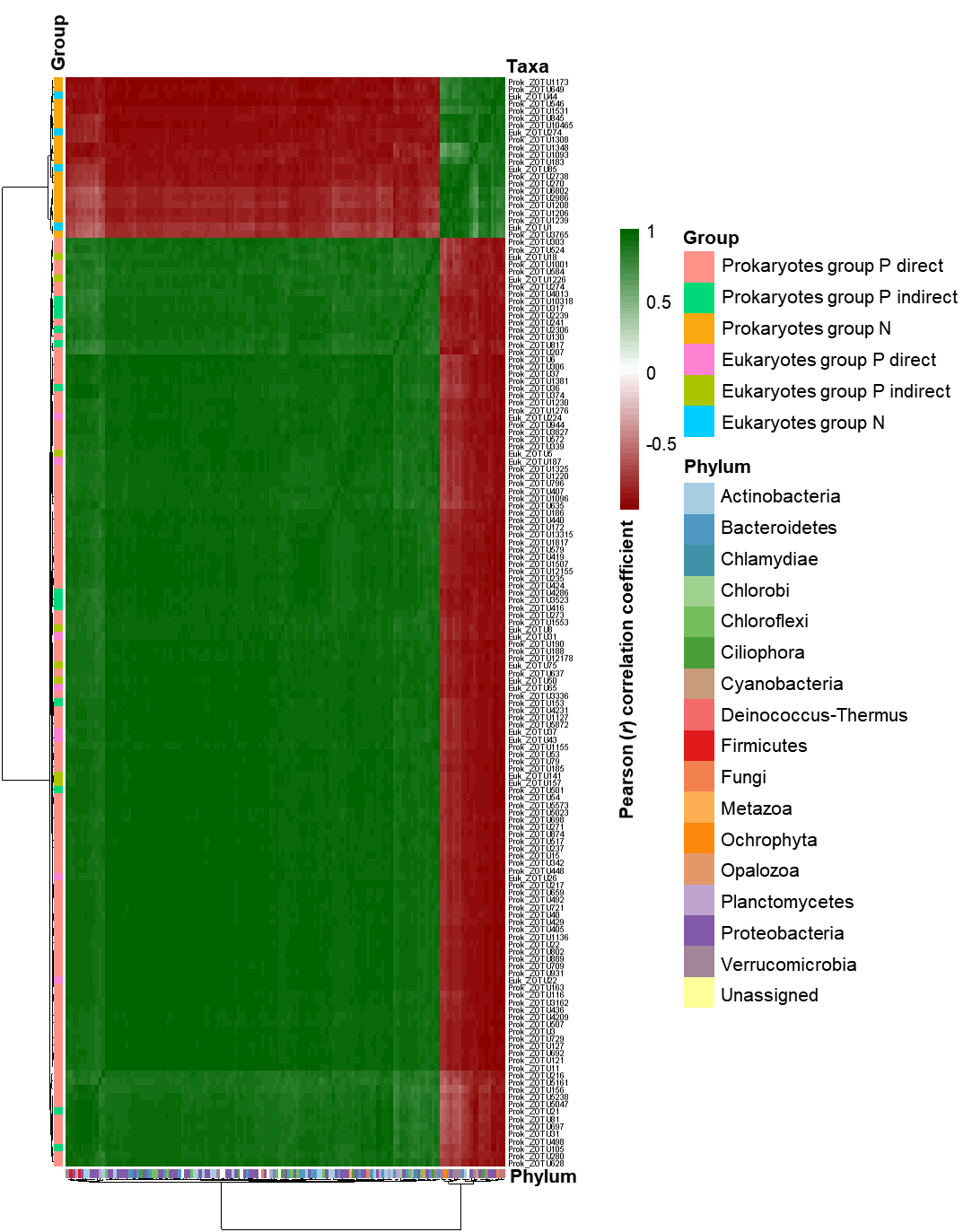

**Figure S6**. Correlation matrix of prokaryotic and eukaryotic taxa selected by the microbial differential abundance testing using DESeq2 and assigned to each group. The groups correspond to *Group P* (*positive*) *direct*, *Group P* (*positive*) *indirect* and *Group N* (*negative*). The heatmap displays the Pearson (*r*) correlation coefficient (*P* < 0.05) based on the relative abundance (variance stabilizing transformation – *vst*-counts) of each taxa in periphyton from all treatments (N = 20). The name of taxa is indicated in each row.

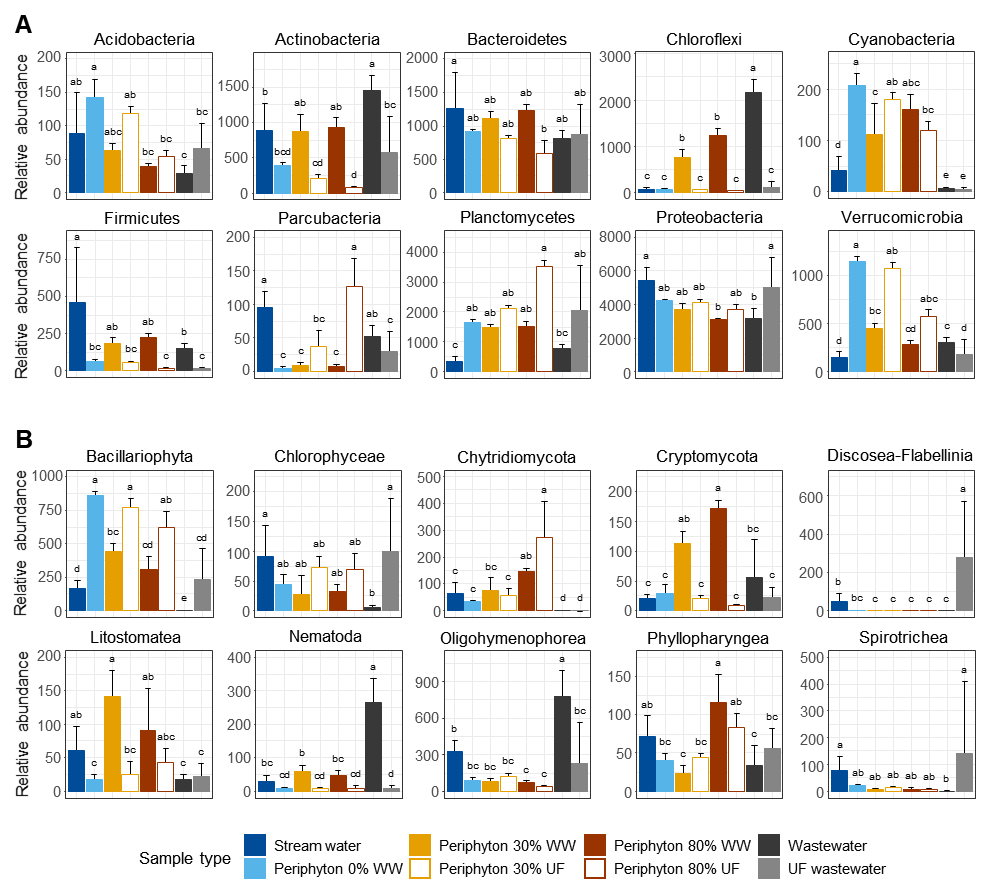

**Figure S7**. Relative abundance of the top-ten prokaryotic phyla (**A**) and eukaryotic classes (**B**) in water samples and in periphyton. The DNA samples were isolated from water samples (stream water, unfiltered wastewater (WW) and ultra-filtered wastewater (UF)) and from periphyton grown in the presence of 0% (control), as well as 30% and 80% WW or UF. Amplicon Sequence Variants (ASVs) abundance for each water (N = 12 sampling times) and periphyton samples (N = 4 channel replicates). Significant differences are indicated by lowercase letters, a > b > c > d > e (Tukey’s test, P < 0.05). Data are mean + SD.
